# Overexpressing NRT2.7 induces nitrate export from the vacuole and increases growth of Arabidopsis

**DOI:** 10.1101/2024.02.26.582130

**Authors:** Patrick Armengaud, Alexis De Angeli, Patrick Berquin, Virginie Bréhaut, Mickaël Durand, Françoise Daniel-Vedele, Anne Krapp, Sophie Filleur

## Abstract

Nitrogen nutrition is essential for crop yield but applying fertilizers has detrimental effects on the environment. Compartmenting nitrate into vacuoles is one of the options to develop Nitrogen-efficient crop adapted to less fertilizers. Only few proteins involved in nitrate transport on the tonoplast have been identified. CLCa is the major transporter involved in nitrate storage in Arabidopsis but it can also facilitate nitrate remobilization from the vacuole in guard cells. Several other nitrate transporters amongst NRT2.7 have been localized in this membrane. The transport mechanism of NRT2.7 has not yet been defined as this protein is present mainly in seed cells that are not easily amenable for electrophysiology analysis. Here, we investigated NRT2.7 function through its ectopic overexpression in a *clca* knock-out mutant. Although the growth diminution of *clca* on nitrogen sufficient medium was complemented, nitrate homeostasis was not restored by NRT2.7 activity. Moreover, NRT2.7 ectopic overexpression in wild-type background (WT) increased growth under limiting nitrogen supply, suggesting that NRT2.7 stimulates nitrate efflux from vacuoles. This hypothesis was demonstrated by electrophysiological nitrate flux measurements on isolated vacuoles. This discovery of NRT2.7 function and more largely the coupling of vacuolar nitrate fluxes with growth under low nitrate supply, will enable new strategies for engineering better NUE for a more sustainable agriculture.

**Highlight:** The overexpression of the nitrate transporter NRT2.7 stimulates growth by increasing the export of nitrate from the vacuole, the main cell compartment for nitrate storage.

## Introduction

Nitrogen (N) is an essential macronutrient acquired by plants from the soil or the air to synthesize amino acids, nucleotides and proteins. Most cultivated plant species rely on soil soluble ammonium and nitrate, the latter being the most abundant oxidized form of nitrogen found in temperate agricultural soils in amounts ranging from 0.1 to 10 mM (Miller *et al*., 2007). The effect of N supply on plant growth has long been evidenced and this nutritional benefit was the basis of the fertilizer-driven yield increases. Crops are therefore abundantly provided with nitrogen fertilizers to insure best yield. This practice is at high economic and ecological costs since it has been shown that plants use only 25 to 50 % of nitrogen applied in the field (Cassman *et al*., 2002). Because of both nitrate mobility and soil structure, nitrate leaks from the field, contaminates water effluents and stimulates growth of algae and microorganisms leading to ecological problems such as river eutrophication (Howarth and Marino, 2006). To avoid these problems and move toward a more sustainable agriculture, one means is to reduce fertilizer inputs without affecting crop yields by improving plant nitrogen use efficiency (NUE). This objective is challenging since NUE is a complex trait that includes uptake, utilization and remobilization efficiencies of the whole plant or of specific organs such as grains (Liu *et al*., 2022). Therefore, research on NUE improvement has been tackled through approaches engineering known molecular players (transporters, enzymes) or heuristically by Quantitative Trait Loci (QTL), Genome-Wide Association Study (GWAS) and transcriptome candidate genes approaches. However, to date, crop varieties with optimized NUE are still rare (Liu *et al*., 2022; Melino *et al*., 2022; Govindasamy *et al*., 2023).

The large number of nitrate transport mechanisms and their multiple level of regulation form a complex network coordinating nitrate fate and homeostasis. The latter is illustrated by remarkably stable cytosolic nitrate levels ranging between 3 and 5 mM (Cookson *et al*., 2005). Once absorbed into root cells, nitrate has three possible fates (Dechorgnat *et al*., 2011). A first one is a direct export to vascular tissue partially through xylem associated nitrate transporters. A second one is its assimilation where cytosolic nitrate is reduced first by the nitrate reductase to nitrite and then by the plastidial nitrite reductase to form ammonium. Ammonium is then incorporated into organic acids to produce amino acids. The third fate of cytosolic nitrate is vacuolar storage. The vacuolar compartment plays important roles in controlling of the plant nitrate status as vacuolar nitrate accumulation has two major functions (Han *et al*., 2016; Hodin *et al*., 2023). First, vacuolar nitrate represents a temporary reserve of N allowing a near constant cytoplasmic provision independently of external fluctuations as nitrate concentrations in this compartment can reach up to 80 mM (van der Leij *et al*., 1998; Cookson *et al*., 2005). Second, vacuolar nitrate acts as an osmoticum contributing to the osmotic potential, including during stomata movements (McIntyre, 1997; Wege *et al*., 2014; Hodin *et al*., 2023).

The first nitrate/proton exchanger mediating nitrate influx into the vacuole, CLCa, was characterized in Arabidopsis (De Angeli *et al*., 2006). This protein is part of a family of seven members in this species. CLCa and CLCb were shown to preferentially exchange nitrate over chloride against proton (De Angeli *et al*., 2006; Bergsdorf *et al*., 2009; von der Fecht-Bartenbach *et al*., 2010; Wege *et al*., 2010; Barbier-Brygoo *et al*., 2011). Both of these exchangers are located on the vacuolar membrane. CLCa activity accounts for up to 50% of vacuolar nitrate content but can also be involved in nitrate efflux from the vacuole in stomata depending of its state of phosphorylation (Geelen *et al*., 2000; Wege *et al*., 2014). Impaired nitrate flux into vacuoles through CLCa was shown to reduce nitrate uptake, nitrate content and expression level of several nitrate transporter genes but to increase nitrate assimilation and consequently NUE (Monachello *et al*., 2009; Hodin *et al*., 2023). As for CLCb, its participation in nitrate accumulation could not been demonstrated as nitrate contents are at the same level in wild type and *clcb* KO mutant when plants are grown in sufficient nitrate-condition (von der Fecht-Bartenbach *et al*., 2010). However, it was proposed recently that CLCb mediates nitrate efflux from the vacuoles when plants are grown on low nitrate concentrations (Shi *et al*., 2023).

Next to CLC transporters, some members of NRT2 and NPF transporters family are localized in the vacuolar membrane (Hu *et al*., 2016; He *et al*., 2017; Wang *et al*., 2018; Lu *et al*., 2022). Among them, NRT2.7 belongs to the seven members of the *NRT2* gene family in Arabidopsis that encode major high affinity nitrate transporters. All NRT2 proteins except NRT2.7 are localized at the plasma membrane. NRT2.7 forms indeed a distinct monophyletic group within the Arabidopsis NRT2 family. The expression of the *NRT2.7* gene occurs mainly in seeds, but its expression has been also described for leaves and roots (Orsel *et al*., 2002; Okamoto *et al*., 2003; Chopin *et al*., 2007). NRT2.7 is able to mediate nitrate fluxes in Xenopus oocytes. The main phenotype of *nrt2*.7 knock-out mutants is a greatly decreased nitrate content in seeds which highlights the major roles of NRT2.7 in nitrate accumulation at reproductive stage (Chopin *et al*., 2007).

Regarding nitrate as a signal, a nutrient or an electro-osmotic charge, further work in characterizing transporters and channels involved in vacuolar fluxes will shed light on the putative sensing and signaling properties of the largest compartment of plant cells. The mechanism of nitrate transport by NRT2.7 was not investigated *in planta* as this protein is essentially present in seeds. So, in this report, we first investigated the impact of a NRT2.7 overexpression in Arabidopsis on nitrate homeostasis, activities of plasma membrane nitrate uptake systems, growth and vacuolar nitrate currents. Functional characterization of nitrate transporters *in planta* often requires the genetic removal of the quantitatively most important components to uncover more subtle or fine-tuning mechanisms. For example, the nitrate transport activity of NRT2 family members are masked by the NRT2.1 activity and has to be assessed in the *nrt2.1* deficient background (Chopin *et al*., 2007; Kiba *et al*., 2012; Lezhneva *et al*., 2014). Here we applied a similar strategy and used the *clca* mutant background in addition to a wild-type background, to test the physiological impact of NTR2.7 activities *in planta* and to determine by electrophysiology the effect the overexpression of NRT2.7 in *clca* on vacuolar nitrate currents. We thus elucidate the function of NRT2.7 by asking the following questions: (i) whether or not overexpression of NRT2.7 nitrate transporter would give similar effects as overexpression of the CLCa nitrate exchanger in the *clca* knock-out genetic background, (ii) how the *NRT2.7* overexpression in wild-type background would affect nitrate homeostasis and growth under nitrate sufficient and limiting conditions and, (iii) what are the consequences of *NRT2.7* overexpression on nitrate fluxes through the vacuolar membrane.

## Material and methods

### Plant material and growth conditions

All *Arabidopsis thaliana* genotypes used in this study were obtained or generated in the Wassilewskija (WS) background. The *clca-2* T-DNA knock-out plant line originates from the Versailles collection (FST 171A06, line EAE89] whereas *clca-1* was isolated in Geelen *et al*. (2000). The *CLCa-OX* (*clca-2*) line (*clca-2* knock-out mutant overexpressing *CLCa* under the 35S promoter of the Cauliflower Mosaic Virus, CaMV) originates from Wege *et al*. (2010). The *NRT2.7-OX* (*clca*) lines (*clca* knock-out mutant overexpressing *NRT2.7* under the 35S promoter of the CaMV) was obtained by crossing *NRT2.7* overexpressors (in the WS background, Chopin *et al*., 2007) with the *clca* mutants (*clca-1* and *clca-2*). The vectors used to over-express *CLCa* and *NRT2.7* contain the three regions necessary for the activity of the *35S* promoter (Ow *et al*., 1987).

Seeds were surface-sterilized, stored for 3 days at 4°C to synchronize germination and sown in 120 x 120 mm square Petri dishes containing either 35 or 70 mL nutrient medium to monitor root architecture (4-5 plants/plate) or to produce material for analysis (12-15 plants/plate). Media differed in their nitrate content (0.2|2|20 mM) and consisted respectively of 0.2|0.7|7 mM KNO_3_; 0|0.65|6.5 mM Ca(NO_3_)_2_; 0.5|0.5|1.5 mM MgSO_4_; 0.625|0.625|0 mM KH_2_PO_4_; 0|0|1.875 mM NaH_2_PO_4_; 0.75|1.25|1.25 mM NaCl; 0.65|0.25|0 mM CaCl_2_ and 1.05|0.55|0 mM KCl. For all media, micronutrients (in µM) consisted of 42.5 FeNaEDTA, 0.16 CuSO_4_, 45 H_3_BO_3_, 0.015 (NH_4_)_6_Mo_7_O_24_, 0.01 CoCl_2_, 0.38 ZnSO_4_, 1.8 MnSO_4_. All media were supplemented with 1% sucrose, pH balanced at 5.8 before autoclaving and solidified using 1% agar (Sigma A1296). Petri dishes were sealed using 3M adhesive tape and placed below tubes delivering 65 µE irradiance for 16h/day in a tissue culture room with a thermoperiod day/night of 22/18°C. At 12 days after germination (14 days after sowing), plants were harvested (or sampled for ^15^N uptake) between 10 and 12 am (after 3 to 5h light exposure).

### Growth measurements

Root architecture was measured at 3, 6 and 9 days after germination by scanning Petri dishes using a flat-bed scanner (Epson Perfection 4490 Photo) at a 200 dots per inch resolution. Total root length was acquired by analyzing root morphology with the EZ-Rhizo software (Armengaud *et al*., 2009). Plant biomass was measured directly on a microbalance (Mettler Toledo XS3DU) for fresh weight.

### RNA extraction and RT-qPCR analysis

RNA was extracted from shoots using the Trizol reagent (Invitrogen) according to the provider recommendations. One microgram was treated with DNase and reverse transcribed using the Quantitect kit (Qiagen) following the manufacturer instructions. For qPCR analysis, specific primers were used for *NRT2.7* (5‘-CCTTCATCCTCGTCCGTTTC-3’ (forward) and 5’-AATTCGGCTATGGTGGAGTA-3’ (reverse)) and *EF1α* ((At5g60390) 5’-CTGGAGGTTTTGAGGCTGGTAT-3’ (forward) and 5’-CCAAGGGTGAAAGCAAGAAGA-3’ (reverse)). The reactions were performed in a Realplex mastercycler (Eppendorf) using commercial SYBR green mastermix (Roche). Samples were subjected to 2 min pre-incubation at 95°C followed by 40 amplification cycles of 15 s at 95°C, 15 s at the melting temperature of the primers set and 20 s at 68°C. *NRT2.7* transcript levels were quantified using linear regression of quantified amplicon range and expressed relatively to *EF1α*.

### Analysis of nitrate, total carbon, and total N contents

Frozen plant material was ground in liquid nitrogen using a Retsch mill grinder (MM301, Retsch GmbH, Germany). For nitrate content determination, an aliquot of 20 mg of frozen powder was used to prepare an ethanol-water extract as described in Cross *et al*. (2006). Nitrate content was assayed after its reduction by vanadium(III) combined with detection by the acidic Griess reaction as described in Miranda *et al*. (2001). Total carbon (C) and N content were determined using an elemental analyzer (EA, Flash 2000, Thermo Scientific) on aliquots of 0.5 to 2 mg dry weight in tin capsules. EA settings consisted of 20 sec of oxygen flux at 80 mL/min and helium at 140 mL/min for 5 min.

### 15N nitrate uptake measurements

Influx of ^15^NO_3_^-^ was assayed as previously described (Orsel *et al*., 2004). Two to three 12-day-old plants were first transferred to 0.1 mM CaSO_4_ for 1 min. To measure high affinity ^15^N nitrate uptake, plants were then transferred to a liquid 0.2 mM nitrate containing medium (the same as described earlier but without sucrose) enriched at 99% with ^15^NO_3_^-^ (Sigma). To measure low affinity transport capacity, plants were instead transferred to a 6 mM nitrate medium consisting of 3 mM K^15^NO_3_; 1.5 mM Ca(^15^NO_3_)_2_; 0.5 mM MgSO_4_; 0.625 mM KH_2_PO_4_; 1.5 mM NaCl and the same micronutrients as described before. ^15^N nitrate uptake was performed for 5 min and plants were finally incubated for 1 min in 0.1 mM CaSO_4_. Roots were then placed into tin capsules, dried at least 48h at 80°C and sent for ^15^N elemental analysis using an ANCA–MS system (PDZ Europa, Crewe, UK) at INRAE Montpellier. Nitrate uptake was calculated from the ^15^N content of the roots (0.5 to 1.5 mg DW). Each uptake experiment included at least 5 measurements per condition and genotype.

### Patch-clamp analyses

Patch-clamp experiments were made in the whole-vacuole configuration with an access resistance between 1.5-3 MΩ. Patch-clamp recordings were made with an EPC8 amplifier (HEKA, Lambrecht-Pfalz Germany). Data were acquired with an AD/DA converter Instrutech LIH 8+8 under the control of the software PatchMaster (HEKA, Lambrecht-Pfalz Germany) and analysed with FitMaster software (HEKA, Lambrecht-Pfalz Germany). Applied membrane potentials were always corrected by the liquid junction potential (Neher, 1992). Current to voltage characteristics were obtained measuring the amplitude of the stationary current. Measurements were done after waiting that the vacuolar lumen solution was equilibrated with the pipette solution, typically 10 minutes after the establishment of the whole-vacuole configuration. Cytosolic side solutions were changed using a gravity driven perfusion system. Bis-Tris-propane (BTP) was chosen as a large impermeable cation in order to minimize cation currents. The solution used for patch-clamp experiments were (in mM): Nitrate vacuolar solution: 100 BTP, 200 HNO_3_, 0.1 CaCl_2_, 5 MgCl_2_, 5 MES, pH = 5.5. Chloride vacuole solution : 475 MES, 15 HCl, 60.5 BTP, 0.1 CaCl_2_, pH = 5.5. Chloride cytosolic solutions: 7.5 BTP, 15 HCl, 0.1 CaCl_2_, 2 MgCl_2_, 15 MES, pH = 7. Nitrate cytosolic solution: 0.1 Ca(NO_3_)_2_, 2 Mg(NO_3_)_2_, 15 MES, pH = 7 with BTP. Osmotic pressures of the cytosolic and vacuolar solutions were adjusted with sorbitol to 640 and 650 mOsm, respectively.

### Statistical analysis

Statistical analyses were conducted using the pairwisePermutationTest function from the rcompanion package in the R environment, except for the electrophysiology results. For those, a Mann-Whitney test was applied using GraphPad Prism 8 software. All the raw data and the results of the statistical analysis can be found in the supplementary table.

## Results

### Both *CLCa* and *NRT2.7* overexpression restore growth of the *clca-2* mutant

To set a suitable genetic background for NRT2.7 functional studies, we characterized the phenotypes of *clca* mutants on a range of nitrate supplied as a sole nitrogen source in vertical Petri dishes. We used *clca-1* and *clca-2* knock-outs (Geelen *et al*., 2000; De Angeli *et al*., 2006) which both showed a significant decrease in shoot fresh weight (-13 to -45%) and root growth rate (-8 to -40%) when compared to the WS wild type (Fig. 1, Supplementary Fig. S1A, B and S2A, B). This decrease was more pronounced under non-limiting nitrate supply (2 and 20 mM). As expected, *clca mutants* showed reduced nitrate content compared to WS under all N supplies in both shoot and root (Supplemental Fig. S1C, D and S2C, D).

**Fig. 1.**
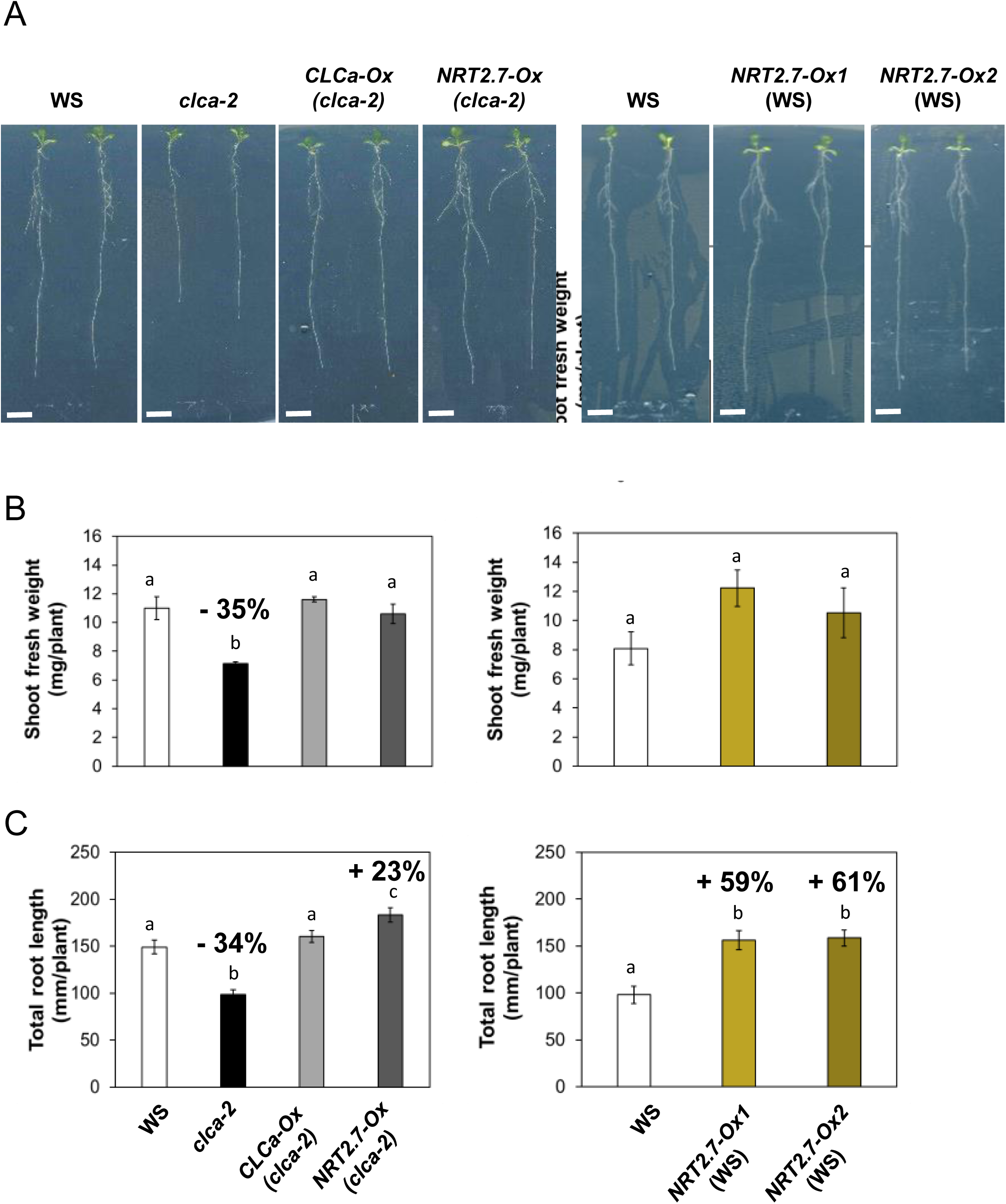
Growth effect of *NRT2.7* overexpression in *clca-2* and wild-type backgrounds. (A) Pictures of wild type (WS), *clca-2*, complemented line (*CLCa-Ox* (*clca-2*)) and *NRT2.7* over-expressing line in *clca-2* (*NRT2.7-Ox* (*clca-2*)) on the left side and, WS and the two *NRT2.7* over-expressing lines, *NRT2.7-Ox1* (WS) and *NRT2.7-Ox2* (WS) on the right side. Plants were grown on 2 mM nitrate for 12 days (Scale bars=1 cm). Shoot fresh weight (B) and total root length (C) were measured on wild type (WS), *clca-2*, *CLCa-Ox* (*clca-2*) and *NRT2.7-Ox* (*clca-2*) (left side of the panel) or WS, *NRT2.7-Ox1* (WS) and *NRT2.7-Ox2* (WS) (right side of the panel). The number of samples were 4-5 for shoot fresh weight (one sample = a pool of 3-4 plants) and n=15-17 for total root length on individual plants. One representative experiment out of three is shown. The numbers above the bars showed the percentage increase or decrease compared to WS. Presented data are means ± Standard Error of the Mean (SEM). Different letters at the top of the bars represent significant difference (p value<0.05).

To study the activity of NRT2.7, we overexpressed ectopically *NRT2.7* in the *clca-2* background and as control in WS. To produce the different genotypes, we used the *NRT2.7*-overexpressing lines *NRT2.7-Ox1 (*WS) and *NRT2.7-Ox2 (*WS) generated in a previous study (Chopin *et al*., 2007) and analysed the *NRT2.7* expression in aerial part. It was very weakly expressed in WS whereas the level of its mRNA is very high in *NRT2.7-Ox1 (*WS) and *NRT2.7-Ox2 (*WS) (Supplementary Fig. S3). These two lines were crossed with the *clca* mutant to obtain *NRT2.7-Ox1* (*clca-2*) and *NRT2.7-Ox2* (*clca-2*), expressing consequently *NRT2.7* at the same level than in the WS background*. NRT2.7-Ox1* (*clca-2*) and *NRT2.7-Ox2* (*clca-2*) showed similar phenotypes and representative data for one line are presented. For comparison, we included in our studies a *clca-2* mutant overexpressing the *CLCa* cDNA under the control of the strong 35S promoter of the CaMV (referred thereafter as *CLCa-Ox* (*clca-2)*) (Wege *et al*., 2010). In this line, *CLCa* transcript levels were increased 30-fold compared to WS (Wege *et al*., 2014).

Based on the more pronounced differences in growth between *clca-2* and WS under non-limiting external N supply, we cultivated the plants on 2 mM nitrate medium for all further analyses using both *clca-2* lines overexpressing either *CLCa* or *NRT2.7* compared to the controls WS, *NRT2.7* over-expressing lines (*NRT2.7-Ox1* (WS)) and *clca-2.* Both *CLCa* and *NRT2.7* overexpression restored the *clca-2* growth phenotypes for shoot biomass and total root length (Fig. 1). This growth restoration was also observed in both lines overexpressing *NRT2.7* in the *clca-1* knock-out background and occurred regardless external nitrate availability (Supplementary Fig. S2A, B). Wild-type plants overexpressing *NRT2.7* showed an important increase of 60% in total root length and no significant difference in shoot fresh weight compared to WT (Fig. 1B, C). The growth stimulation observed for *NRT2.7* overexpressing lines both in the *clca-2* and the WS background, is therefore independent of the *clca-2* mutation. The physiological changes triggered by *NRT2.7* overexpression in the *clca* mutants are thus different from the complementation by *CLCa* while both led to restored growth.

### *NRT2.7* and *CLCa* overexpression have opposite effects on both nitrate content and nitrate uptake capacities

To decipher the underlying mechanism that impact growth of these overexpressing lines, we determined nitrate content in roots and shoots. As described earlier (Wege *et al*., 2010), nitrate content of *clca-2* was 35 and 28 % lower than WT level in shoots and roots, respectively (Fig. 2A, C). Expression of *CLCa* in the *clca-2* mutant restored nitrate levels in shoots and roots, whereas *NRT2.7* overexpression in the *clca-2* background led to a 53% decrease of the nitrate content when compared with the *clca-2* mutant (Fig. 2A, C). Likewise, *NRT2.7* overexpression does not complement *clca-1* phenotype for shoot and root nitrate content regardless the external nitrate concentration for plant growth (Supplementary Fig. S2C, D). This showed that the increased growth observed in the *NRT2.7-Ox (clca-2)* and *NRT2.7-Ox (clca-1)* lines occurred without restoring nitrate content. Like the effect of *NRT2.7* overexpression in the *clca-2* mutant background, overexpression of *NRT2.7* in wild-type background was accompanied by a slight decrease in nitrate content in roots but no significant modification was observed in shoots (Fig. 2B, D).

**Fig. 2.**
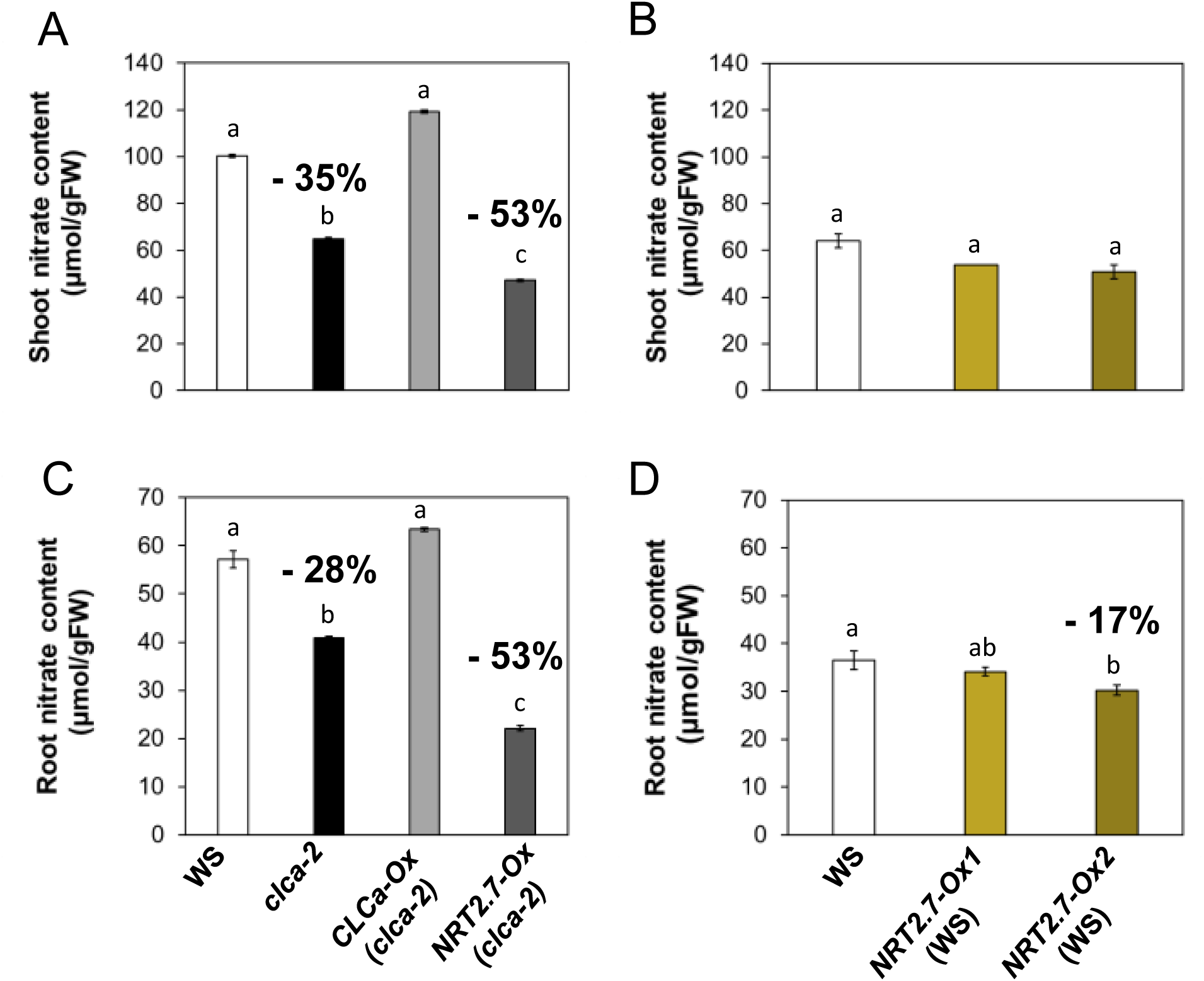
Nitrate content of *NRT2.7* overexpression in *clca-2* and wild-type backgrounds. Wild type (WS), *clca-2*, complemented line (*CLCa-Ox* (*clca-2*)) and *NRT2.7* over-expressing line in *clca-2* (*NRT2.7-Ox* (*clca-2*)) (A, C) or WS and the two *NRT2.7* over-expressing lines, *NRT2.7-Ox1* (WS) and *NRT2.7-Ox2* (WS) (B, D), were grown on 2 mM nitrate. After 12 days on the medium, shoot (A, B) and root (C, D) nitrate contents were measured. Two to three biological replicates were performed on each genotype, one sample corresponding to a pool of at least 50 plants. One representative experiment out of two is shown. The numbers above the bars showed the percentage increase or decrease compared to WS. Presented data are means ± SEM. Different letters at the top of the bars represent significant difference (p value<0.05).

In order to understand the impact on nitrate levels, we investigated how nitrate uptake was affected in these lines. Nitrate uptake was measured using ^15^N labelled nitrate at concentration of 0.2 and 6 mM to monitor both high and low affinity transport system activities (HATS and LATS, respectively, this latter activity being obtained as the difference between uptake measurements at 6 mM and 0.2 mM). Figure 3 shows that the HATS activity was decreased by 16% in *clca-2* whereas the LATS activity was not significantly different from that of WT. The *CLCa-Ox* (*clca-2*) line exhibited the same uptake capabilities for both transport activities as wild-type plants, illustrating a full complementation of the *clca-2* mutant phenotype. The *NRT2.7-Ox* (*clca-2*) line showed similar HATS activity as the *clca-2* knock-out mutant but an increase by 29% of LATS activity (Fig. 3B, C). The *NRT2.7-Ox* (WS) lines displayed comparable profiles but enhanced effect with a 2-fold increased activity of the LATS when compared to WT (Fig. 3C). Thus, whereas *CLCa* overexpression in the *clca* mutants increased mainly HATS activity compared to WT, *NRT2.7* overexpression had the opposite effect, supporting the hypothesis that these two nitrate transporters have different functions *in planta* when overexpressed ectopically.

**Fig. 3.**
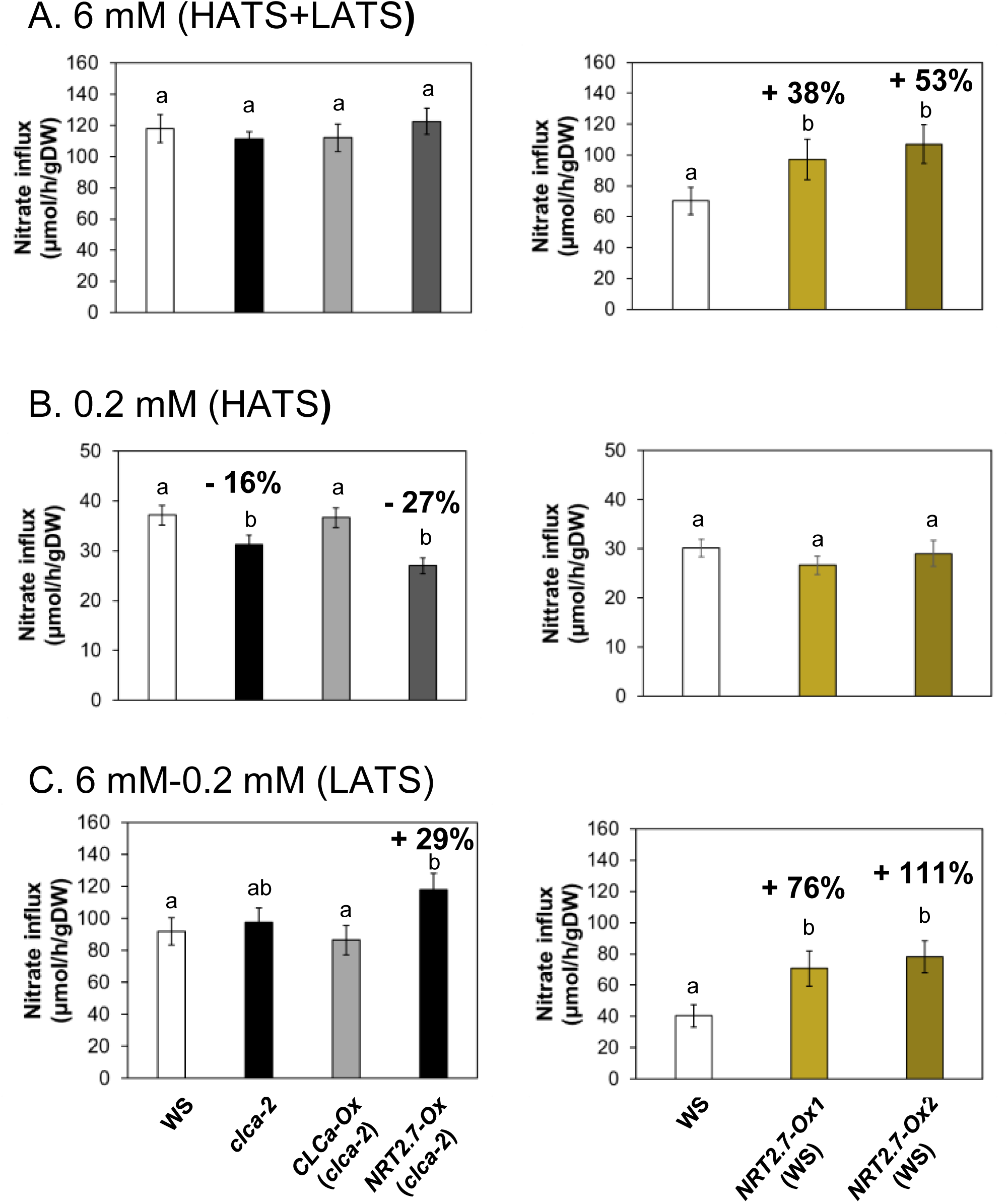
Effect of *NRT2.7* overexpression in *clca-2* and wild-type backgrounds on nitrate uptake activities. Wild type (WS), *clca-2*, complemented line (*CLCa-Ox* (*clca-2*))*, NRT2.7* over-expressing line in *clca-2* (*NRT2.7-Ox* (*clca-2*)) and the two *NRT2.7* over-expressing lines, *NRT2.7-Ox1* (WS) and *NRT2.7-Ox2* (WS), were grown on vertical Petri dishes on a medium containing 2 mM nitrate for 12 days. Uptake experiments consisted of four to five measurements of ^15^NO_3_^-^ in roots from seedling that were incubated for 5 min in a liquid medium containing 6 mM (A = HATS+LATS) or 0.2 mM (B = HATS) ^15^NO_3_^-^ . The activity of LATS (C) was obtained by subtracting the activity measured at 0.2 mM to the one at 6 mM (see material and methods section for details). Presented data are averages of three independent experiments (n=3-5 by independent experiment). Bars represents means ± SEM. The numbers above the bars showed the percentage increase or decrease compared to WS. Different letters at the top of the bars represent significant difference (p value<0.05).

### *NRT2.7* overexpression leads to increased biomass under limiting and non-limiting conditions

Plants stimulate their root growth when their N nutritional status is low (Jia and von Wirén, 2020). Given the important increase of root growth in the *NRT2.7-Ox* (WS) lines under non-limiting N supply (Fig. 1), we further studied the growth-promoting effect of *NRT2.7* overexpression under limiting N supply to better understand the impacts of the changes in nitrate fluxes through the vacuolar membrane. Plants were grown on vertical agar medium containing 0.2 mM nitrate for 12 days. In these conditions, shoot biomass and total root length of *NRT2.7-Ox* (WS) lines were increased when compared to WT by 49 to 65% and 19 to 33% respectively (Fig. 4A, B). Conversely, root nitrate content of the *NRT2.7-Ox* (WS) lines was dramatically reduced by 75 to 83% compared to WT, whereas shoot nitrate content was not modified (Fig. 4C, D). A similar tendency was observed when *NRT2.7* was overexpressed in *clca* KO mutant comparing nitrate content to shoot and root biomass (Supplementary Fig. S4).

**Fig. 4.**
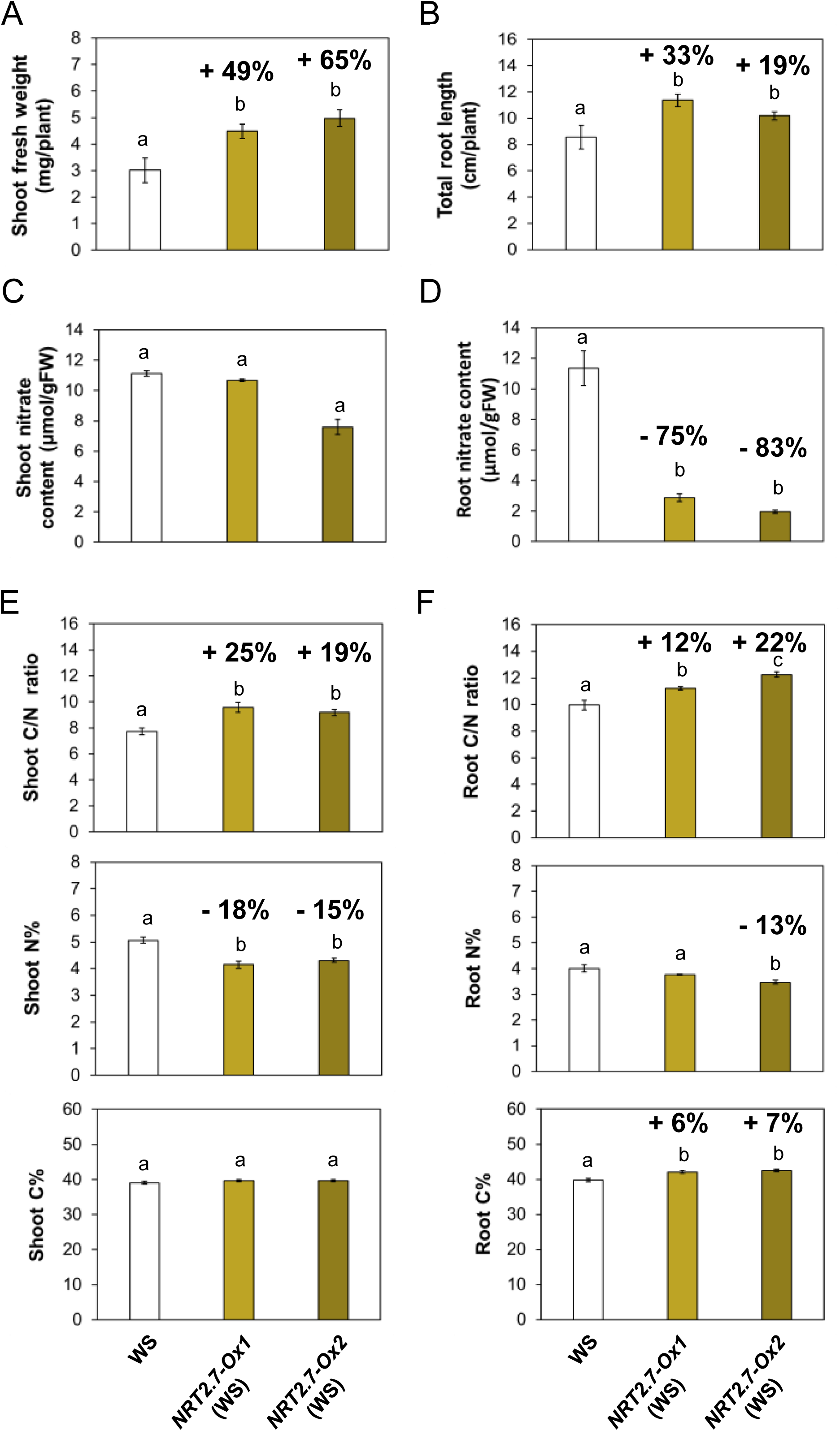
Effect of limiting N supply on the phenotype of *NRT2.7* over-expressers. Wild type (WS) and over-expressing lines, *NRT2.7-Ox1* (WS) and *NRT2.7-Ox2* (WS), were grown in petri dishes for 12 days on 0.2 mM nitrate. Shoot fresh weight (A), total root length (B) shoot and root nitrate contents (C, D) were measured as described in Figure 1 and 2. C/N ratios, percentages of N and C contents were measured in shoot (E) and root (F) using an elementary analyser. Data are averages of four biological replicates for shoot fresh weight, 12 to 17 for total root length, two to four for nitrate contents and four for C and N contents. The numbers above the bars showed the percentage increase or decrease compared to WS. Different letters show significant differences between genotypes (p<0.05).

The carbon to nitrogen (C/N) ratio is an essential indicator that plants perceive to orientate their growth, development and metabolism (Coruzzi and Zhou, 2001). The C/N ratio of *NRT2.7-Ox* (WS) compared to WT was increased in shoots and roots mainly due to a decrease in total N proportion (Fig. 4E). These data suggest that the ectopic expression of *NRT2.7* drives nitrate use for biomass production. As no difference in nitrate uptake through the HATS was observed (Fig. 3), the strong decrease in nitrate content could be the consequence of a lower vacuolar storage capacity and/or an increased nitrate flux out of the vacuole as it was previously shown for other vacuolar nitrate transporters (He *et al*., 2017; Lu *et al*., 2022; Hodin *et al*., 2023; Shi *et al*., 2023).

### *NRT2.7* overexpression leads to increased nitrate fluxes out of the vacuole

In order to associate the observed physiological changes with NRT2.7 functional properties, we used an electrophysiological approach to determine if the over-expression of *NRT2.7* modifies the nitrate fluxes across the vacuolar membrane. We took advantage of the *NRT2.7-Ox* (*clca-2*) line where such modification would not be masked by CLCa-driven fluxes. We extracted vacuoles from mesophyll protoplasts of the *clca-2* and the two *NRT2.7-Ox* lines (*clca-2*) and performed patch clamp experiments in whole-vacuole configuration (Fig. 5). In order to maximize the fluxes of nitrate directed from the vacuole to the cytosol and from the cytosol to the vacuole, we performed two independent series of experimentations designed as follow. In both experiment series, we used the non-permeant cation BisTriPropane in order to limit the vacuolar currents mediated by cations. With the aim to highlight the efflux of nitrate from the vacuole (positive ionic currents), in the first series of experiments (Fig. 5B), we used bi-ionic conditions with 200 mM nitrate in the vacuole and 19.2 mM chloride in the cytosolic-side solution. The vacuoles extracted from *clca-2* knock-outs presented a mean current density of +1.3 ± 0.4 pA/pF at +43 mV and a mean reversal potential (E_rev_=-8 ± 3 mV) that were comparable to those previously observed in similar ionic conditions (De Angeli *et al*., 2006; Wege *et al*., 2010). Importantly, in the same ionic conditions, vacuoles from *NRT2.7-Ox (clca-2)* plants presented significantly higher current densities compared to *clca-2* currents only at positive vacuolar membrane potentials (from +3 mV to +83 mV, Fig. 5B). To note, similar results were obtained for both lines as displayed on the figures by vertical or horizontal diamonds. These positive currents are compatible with an increased nitrate efflux capacity of the vacuoles from *NRT2.7-Ox (clca-2)* plants. In the second set of experiments, we used an ionic configuration suited to highlight nitrate currents directed from the cytosol to the vacuolar lumen. We performed cytosolic-side solution exchange experiments starting from a chloride-based buffer (19.2 mM Cl^-^ pH 7) to a nitrate based buffer (4 mM NO_3_^-^ pH 7) keeping the vacuolar-side buffer constant and with only chloride as a permeant anion (15 mM Cl^-^ pH 5.5) (Supplementary Fig. S5A and S5B). Upon the exchange of the cytosolic-side solution a similar increase of the negative current was observed in vacuoles obtained from the tested genotypes. Indeed, the mean current ratios at -60 mV were I_NO3_/I_Cl_=1.84±0.4 in *NRT2.7-Ox (clca-2)* vacuoles and I_NO3_/I_Cl_=1.77 ± 0.44 in *clca-2* vacuole*s*. Conversely, the current densities in presence of nitrate did not show a significant difference between the genotypes, being I_NO3_=-2.4±1 pA/pF at -60 mV in *NRT2.7-Ox* (*clca-2*) and I_NO3_=-2.7±0.9 pA/pF at -60 mV in *clca-2.* As the current amplitudes in *NRT2.7* overexpressor show a high variability compared to those of the *clca-2* mutant, we checked whether the rectification ratios between the currents at +83 mV versus -77 mV and +63 mV versus -57 mV (Supplementary Fig. S5C and S5D) were not different between the vacuoles from the different genotypes. The values of these ratios were around -3 and there was no significant difference between the two genotypes. Further no correlation between the rectification ratio (I_+83mV_/I_-77mV_) and current density (I_+83mV_) could be identified. This indicates that the high variability of currents obtained in *NRT2.7*-Ox (*clca-2*) is not linked to background leak currents. In conclusion, this second series of experiments indicates that the vacuoles obtained from *NRT2.7-Ox* (*clca-2*) do not display an increased capacity of driving NO_3_^-^ fluxes directed to the vacuolar lumen compared to the vacuoles from *clca-2* under our experimental conditions. Altogether, the results obtained by patch-clamp experiments suggest that, compared to vacuoles from *clca-2*, *NRT2.7-Ox* (*clca-2*) plant vacuoles present a higher capacity to drive nitrate fluxes directed into the cytosol but are not modified concerning nitrate influx into the vacuole.

**Fig. 5.**
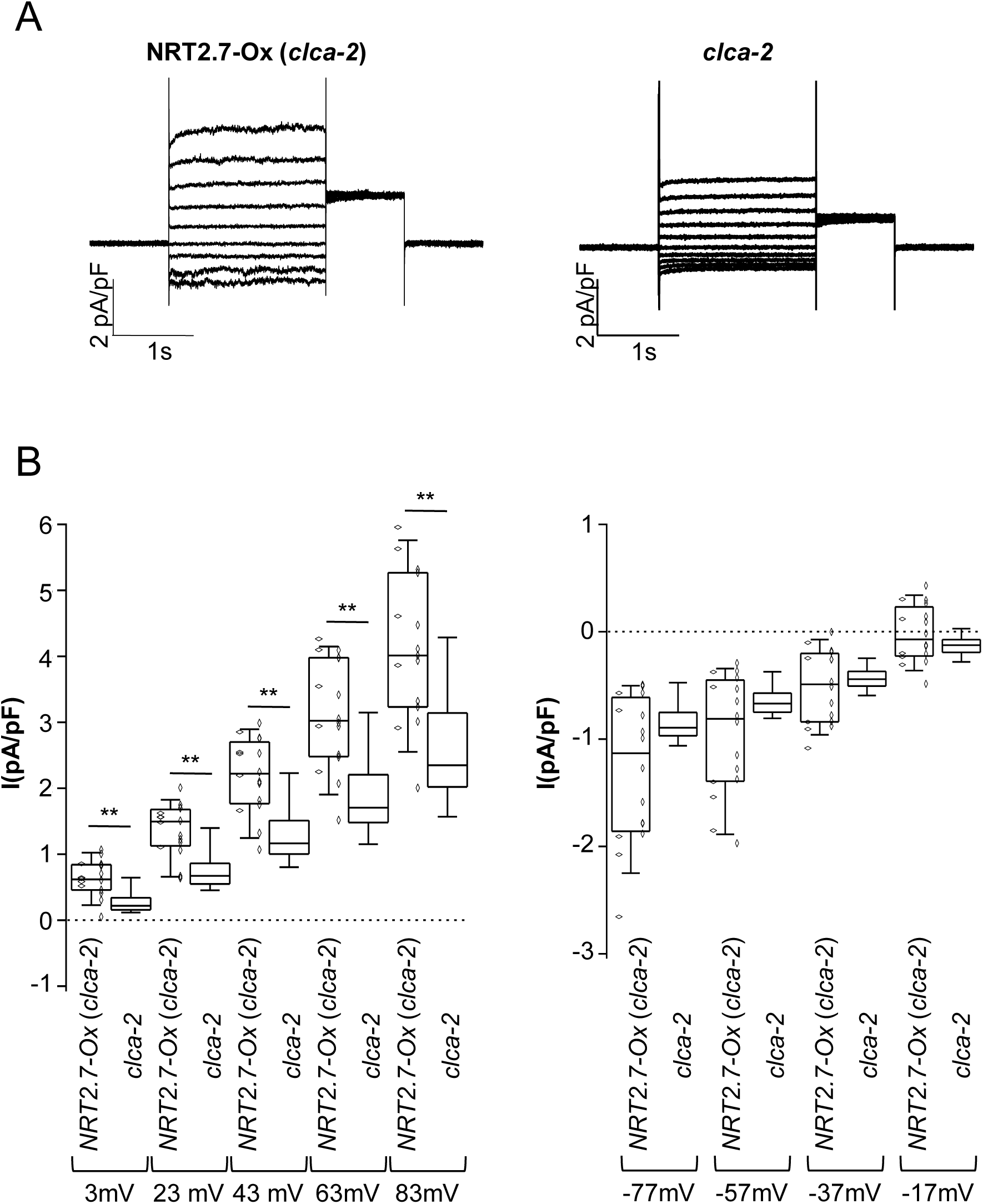
Nitrate efflux from mesophyll vacuoles is increased in *NRT2.7-Ox* (*clca-2*) plants compared to *clca-2* knock-outs. (A) Representative whole-vacuole currents measured in vacuoles extracted from *clca-2* knock-out mutant (left) and *NRT2.7-Ox* (*clca-2*) transgenic line (right) in presence of nitrate only in the vacuolar lumen (200 mM NO_3_^-^ pH 5.5 in the vacuolar buffer, 19.2 mM Cl^-^ in the cytosolic buffer, see Materials and Methods). Currents were evoked in response to 2 s voltage pulses ranging from -77 mV to +83 mV in +20 mV steps, followed by a tail pulse at +33 mV, with a holding potential at -17 mV. (B) Box plot of the mean current density (pA/pF) measured at different positive (left) and negative (right) membrane potentials in vacuoles from *NRT2.7-Ox* (*clca-2*) (vertical diamonds line *NRT2.7-Ox1* (*clca-2*), n=11; horizontal diamonds line *NRT2.7-Ox2* (*clca-2*), n=5), and *clca-2* knock-out plants, n=10. The ionic conditions were as in (A). Boxes defines the first and the third quartile and the median, the whiskers represent the ninth and the ninety-first percentiles. The two asterisks indicate a statistical significance at 1% level with a Mann-Whitney test.

## Discussion

Recent studies underline the importance of nitrate fluxes through the vacuolar membrane to enhance NUE by increasing efflux of nitrate from vacuole and/or reducing nitrate storage in this latter compartment (Han *et al*., 2016; Wang *et al*., 2018; Hodin *et al*., 2023; Shi *et al*., 2023). It is therefore essential to characterize the different vacuolar nitrate transporters and their activities to get a better understanding of their functional specificities. In this report, we analyzed the function of the vacuolar NRT2.7 transporter by introducing its cDNA under the control of the *35S* promoter in *clca* KO mutant or in wild type. The combined results of the electrophysiological analyses and physiological traits of the *CLCa* and *NRT2.7* overexpressing lines strikingly demonstrate that both NRT2.7 and CLCa impact N use but through clearly distinct mechanisms. The *NRT2.7* overexpression in the *clca-2* mutant bypassed the CLCa-dependent reduced growth via a mechanism that uncoupled growth and nitrate accumulation probably related to modified nitrate efflux from the vacuole. This mechanism, when acting in a wild-type background overexpressing *NRT2.7*, led to an increase of biomass production mainly through a higher root biomass.

### NRT2.7 stimulates nitrate export out of the vacuole when overexpressed ectopically

In this study, we used the *CLCa* mutant altered physiology to assess functions and physiological consequences of *NRT2.7* ectopic expression. We chose CLCa over its closest homologous, CLCb, as the *clca* KO mutant has a stronger phenotype than *clcb* in respect to nitrate contents (Geelen *et al*., 2000; von der Fecht-Bartenbach *et al*., 2010; Shi *et al*., 2023) and, nitrate currents are very small in *clca* even when *CLCb* is over-expressed into this mutant compared to wild type (De Angeli *et al*., 2006; Monachello *et al*., 2009). Furthermore, similar to CLCb, CLCa is involved in nitrate efflux out the vacuole depending on its phosphorylation state (Wege *et al*., 2014). As the phosphorylation site is conserved in CLCb, we can assume that CLCb could work in both ways depending on the growth conditions.

In this study, we confirmed that *CLCa* mutant physiology was deeply affected by its limiting capacity of loading nitrate into vacuoles (Fig. 1 and 2; Supplementary Fig. S1 and S2). Besides CLCa, the NRT2.7 nitrate transporter is located at the tonoplast and has been showed to translocate nitrate in Xenopus oocytes. Its loss-of-function reduces nitrate accumulation in seeds where the gene is chiefly expressed (Chopin *et al*., 2007). Although *NRT2.7* mRNA were detected in leaves and roots, no phenotype other than seed traits have been reported in the KO mutant (Orsel *et al*., 2002; Okamoto *et al*., 2003; Chopin *et al*., 2007).

To better characterize its function *in planta*, we overexpressed *NRT2.7* in wild type and *clca* KO mutant backgrounds. The overexpression of *NRT2.7* fully restores the reduced *clca-2* plant growth and even exceeds wild-type biomass but, surprisingly, further diminished the low *clca-2* nitrate content (Fig. 1 and 2). It led to 35% and 23% decreases of shoot and root nitrate pools, respectively, when compared to *clca-2*. These results showed that *NRT2.7* could indeed rescue the *clca-2* growth defect but rather through an alternative mechanism than a true functional complementation. This hypothesis was confirmed as the *NRT2.7* overexpressors in both wild-type and *clca-2* background showed an increase in total root length and a depletion of root nitrate content compared to wild-type plants.

The decrease of root nitrate content caused by *NRT2.7* ectopic overexpression is difficult to conceive with a vacuolar nitrate importer. Chopin *et al*. (2007) reported an influx of ^15^N labeled nitrate into NRT2.7 expressing oocytes. This result indicates that, in this biological system, NRT2.7 was targeted to the plasma membrane with presumably its cytoplasmic N- and C-terminal domains oriented to the oocyte cytoplasm. *In planta*, NRT2.7 is localized at the tonoplast (Chopin *et al*., 2007). As NRT2.7 cytoplasmic terminus is probably maintained in the plant cell cytoplasm (Jacquot *et al*., 2020), NRT2.7 would thus function as a nitrate efflux system from the vacuole to the cytoplasm. This orientation is in agreement with the fact that NRT2s are H^+^/NO_3_^-^ symporters (Forde and Clarkson, 1999) and that roots of *NRT2.7* overexpressors become depleted of nitrate at low external nitrate availability (Fig. 4, Supplementary Fig. S4). In addition, the determination of nitrate fluxes by patch clamp techniques showed that isolated vacuoles from *NRT2.7-Ox* (*clca-2*) plants displayed a higher capacity to drive nitrate fluxes directed to the cytosol th an isolated vacuoles from *clca-2* plants (Fig. 5). The difference of current density between the two genotypes is rather small (i.e about 1.3 pA/pF more in *NRT2.7-Ox* (*clca-2*) vacuoles at +63 mV). This can be explained by a difference of electrogenicity and unitary turnover rate between the two kinds of ion transporters, CLCa being a nitrate/proton antiporter and NRT2.7 a nitrate/proton symporter (Forde and Clarkson, 1999; De Angeli *et al*., 2006). In the same experimental setup, we showed that the capacity of such isolated vacuoles to drive nitrate influx did not vary between these genotypes (Fig. 5 and Supplementary Fig. S5). This indicates that NRT2.7 stimulates vacuolar nitrate export when ectopically overexpressed in seedlings and could explain the observed decrease in nitrate content when *NRT2.7* is overexpressed in *clca-2* background. As this latter effect was also detected in *NRT2.7-Ox* (WS) mainly when the plants were grown on 0.2 mM nitrate (Fig. 4 and Supplementary Fig. S4), the activity of NRT2.7 to stimulate nitrate export does not seem to rely on the import activity of CLCa. Additionally, based on these experiments, we cannot completely exclude that NRT2.7 may regulate positively the activity of nitrate exporters localized in the tonoplast and is not itself the active transporter driving the fluxes. *CLCb*, recently shown to be a nitrate exporter when nitrate is low in the external medium and, overexpressed in *clca* KO mutant (Monachello *et al*., 2009; Shi *et al*., 2023), could have its activity induced by *NRT2.7* overexpression. However, the decrease of root nitrate content by -75 and -83% in *NRT2.7-Ox* (WS) compared to wild type was accompanied by a total root growth increase of 33% when plants are grown on 0.2 mM nitrate (Fig. 4). Under the same growth conditions, a similar tendency was observed in *NRT2.7-Ox* (*clca2),* a 19% increase in root length was accompanied by a 84% reduction in root nitrate content (Supplementary Fig. S4). As *CLCb* is over-expressed in *clca* mutant (Monachello *et al*., 2009), this that CLCb might not be involved in the stimulation of nitrate export by *NRT2.7* overexpression.

Chopin *et al*. (2007) observed a decrease in seed nitrate content in *nrt2.7* mutants, which was interpreted as a reduced capacity to import nitrate into the vacuole of the embryo. *NRT2.7* is highly expressed in seeds and the *NRT2.7* promoter drives expression in the embryos where the protein is located at the tonoplast. A decreased nitrate content is thus difficult to explain by the loss-of-function of a vacuolar nitrate exporter protein. However, the transport direction of NRT2.7 in seeds and in shoot/root vacuoles might be different because of distinct types of vacuoles existing in these organs (Shimada *et al*., 2018). In addition, tonoplast membrane potential might vary and the transport direction may reflect the organ specific ionic and osmotic conditions as it has been discussed previously for NPF6.3 (NRT1.1/CHL1, (Léran *et al*., 2014) or CLCa (Wege *et al*., 2014). Besides, nitrate transport properties are regulated via posttranslational protein modifications (e.g. Wege *et al*., 2014; Jacquot *et al*., 2017), a mechanism that should be explored in the future for NRT2.7. However, no phosphorylation site in NRT2.7 is found in the PhosPhAt databases (Heazlewood *et al*., 2008). Finally, another hypothesis would be that *nrt2.7* seed nitrate content is decreased because nitrate transport from the roots or shoots towards the seeds is diminished in the *nrt2.7* mutants. Indeed, in Arabidopsis seeds, nitrate content seems to rely directly on the nitrate taken up by roots (Schulze *et al*., 1994). Inflorescence grafting experiments are required to distinguish between seed prone or root/shoot prone determination of the decreased nitrate level in *nrt2.7* seeds. Altogether, these results support that NRT2.7 is essential for nitrate fluxes through the vacuolar membrane. It stimulates nitrate export from the vacuole in seedlings but the contrasting phenotype in reproductive and vegetative organs when the expression of *NRT2.7* is modified, poses a challenging hypothesis and needs further investigation.

### Nitrate fluxes through CLCa and NRT2.7 alter root nitrate uptake

As previously reported, we confirmed that the mutation in *CLCa* reduced nitrate uptake activity in the high affinity range (Monachello *et al*., 2009, Fig. 3) whereas the level of LATS activity was not modified. These results illustrate the existence of a coordinated signaling crosstalk between vacuolar and plasma membrane nitrate fluxes. An obvious signal for this coordinated transmembrane transport would be the nitrate anion itself. It was shown that loss of function of CLCa leads to a faster increase of the cytosolic nitrate concentration in response to an increase of the extracellular NO_3_^-^ concentration in guard cells (Demes *et al*., 2020). In this scenario, a transitory elevated cytosolic nitrate level would decrease nitrate uptake, increase the nitrate efflux from the cells and/or the assimilation pathways through the nitrate reductase (NR). This was confirmed in a recent study demonstrating that the inhibition of nitrate storage in the vacuole leads to an increase of the NR activity and a better NUE (Hodin *et al*., 2023). In addition to nitrate, nutrient uptake activity is controlled by a wide range of biophysical (membrane potential, pH) or physiological signals (phosphorylation, metabolites) (see Amtmann and Blatt, 2009). CLCa through its activity as a NO_3_^-^/H^+^ exchanger is known for its function in adjusting the cytosolic pH in guard cells (Demes *et al*., 2020). More recently, Jain and Schmidt (2024) showed that NRT2.1 phosphorylation level varies according to the environmental pH certainly to adjust cytosolic pH. Then the loss of function of *CLCa* could lead to the inactivation of NRT2.1 to compensate the change in cytosolic pH.

The increase of nitrate export from the vacuole due to the *NRT2.7* overexpression leads probably to change in nitrate cytosolic content and/or cytosolic pH, as for *clca-2*. Even if this hypothesis needs to be tested using the nitrate/pH biosensor, ClopHensor (Demes *et al*., 2020), these modifications could explain the decrease in HATS activity observed in *NRT2.7-Ox (clca-2)* like in *clca-2* plants but not the increase of the LATS activity (Fig. 3). This latter discrepancy could suggest that alterations in nitrate influx and efflux in and out of the vacuole do not have the same effect on nitrate uptake. The coordination between the direction of vacuolar nitrate fluxes and nitrate influx at the plasma membrane will need to be further studied in the future.

### *NRT2.7* overexpression increases growth at low external and internal nitrate

The overexpression of *NRT2.7* stimulates nitrate export from the vacuole, leading to an increase in plant growth. This phenotype could be explained not only by the increase of LATS activity but also by a higher nitrate long-distance transport from root to shoot. A study characterizing two *Brassica napus* genotypes differing by their capacity in nitrate storage in root previously showed that the decrease of root nitrate content induces a higher nitrate translocation from root to shoot leading to a better NUE (Han *et al*., 2016). We can assume that a similar phenomenon occurred in the *NRT2.7* overexpressors.

Interestingly, both shoot and root biomass increased in *NRT2.7-Ox* (WS) grown under limiting N nutrition even that a severe decrease of nitrate in root (-75 to -83% compared to wild type) and a slight decrease of total nitrogen content (-8 to –19% compared to wild type) occurred in these growth conditions (Fig. 4). These results confirm an apparent disconnection between growth and nitrate content in *NRT2.7* overexpressors plants. C and N metabolism are interconnected and regulate each other (Zheng, 2009). Interestingly, the C/N ratio was higher in the overexpressors compared to wild type due to a decrease in N proportion indicating that, even if plants are grown under low external N availability, the overexpressors are able to maintain C proportions by regulating photosynthesis activity facilitating the increase in biomass. The low internal nitrate levels could be then the consequence of N dilution occurring under accelerated growth. At the opposite, the increase of nitrate export from the vacuole could stimulate nitrate assimilation and consequently photosynthesis as shown in a *Brassica napus* genotype with reduced root nitrate storage (Han *et al*., 2016) or, more recently when Arabidopsis plants overexpressing the CLCb nitrate vacuolar exporter are grown on low nitrate (Shi *et al*., 2023).

In conclusion, we show that the two vacuolar nitrate transporters CLCa and NRT2.7 share limiting functionality and are not exchangeable. Our results suggest that NRT2.7 stimulates vacuolar nitrate export when overexpressed ectopically. By exploiting genotypes with modified levels of vacuolar transporters, we show that changes in the nitrate influx or efflux from the vacuolar compartment lead to differences in root nitrate uptake suggesting the existence of a fine regulation between vacuolar and plasma membranes through biophysical and physiological networks of vacuolar nitrate sensing and homeostasis. NRT2.7, whose function *in vivo* seemed to be restricted to seeds, triggers new physiological features when ectopically overexpressed. The link between NRT2.7-driven nitrate fluxes and C content requires further studies but may be the basis for improving plants with higher C/N ratios accompanied by increased growth when N is limiting.

## Supplementary data

The following supplementary data are available at *JXB* online.

Fig. S1. Growth and nitrate content of *clca-2* knock-out mutant grown on different nitrate concentrations.

Fig. S2. Growth and nitrate content of *NRT2.7* overexpressors in the c*lca-1* knock-out mutant.

Fig. S3. Level of NRT2.7 expression in *NRT2.7-Ox* (WS) lines.

Fig. S4. Effect of limiting N supply on the phenotype of *NRT2.7* over-expressors in *clca-2* background.

Fig. S5. Nitrate influx in vacuoles from *NRT2.7*-Ox (*clca-2*) plants and from *clca-2* knock-out.

## Acknowledgements

The authors are very grateful to Pascal Tillard for servicing the ^15^N measurements, Anne Marmagne for helping with the elementary C/N analyzer and the IJPB platform ‘The Plant Observatory’, in particular Joël Talbotec and Philippe Maréchal for taking care of greenhouse cultures.

## Author contributions

FDV, AK and SF : conceptualization design ; PA, FDV, ADA, AK and SF: experimental design ; PA, ADA, PB and VB performed the experimental work ; PA, MD, FDV, ADA, AK and SF analyzed the data; PA, FDV, AK and SF wrote the manuscript. All authors reviewed the manuscript.

## Conflict of interest

The authors declare no conflict of interest.

## Funding

This work benefits from the support of the LabEx Saclay Plant Sciences-SPS (ANR-10-LABX-0040-SPS) and of the Agence Nationale de la Recherche (ANR-Nitrapool ANR08-Blan-008).

## Data availability

All the raw data supporting the findings of this work are available upon request from the corresponding authors, Anne Krapp and Sophie Filleur. Statistical analyses are available in the supplementary table 1.

## Abbreviations

CaMV: Cauliflower Mosaic Virus
CLC: ChLoride Channel
KO: Knock-out
N: Nitrogen
NPF: Nitrate transporter 1/Peptide transporter Family
NRT2: High affinity Nitrate transporter
NUE: Nitrogen Use Efficiency
WS: Wassilewskija Arabidopsis accession

---

**Supplementary Fig. S1.**
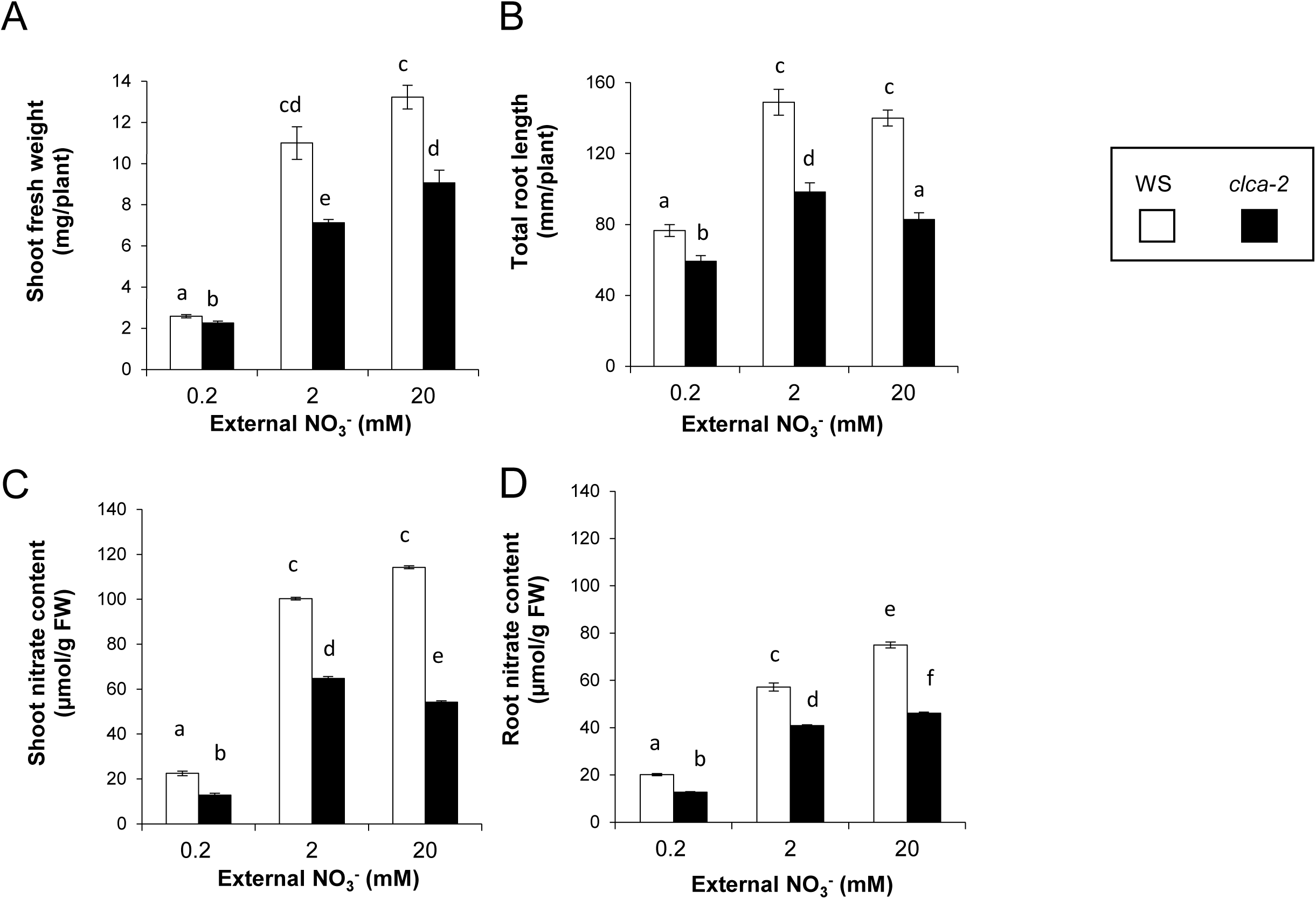
*clca-2* knock-out mutant phenotype. Wild-type (WS) and *clca-2 p*lants were grown on vertical Petri dishes for 12 days on 0.2, 2 or 20 mM external nitrate as described in the ‘Materials and Methods’. Shoot fresh weight (A), total root length (B), shoot and root nitrate contents (C and D) were measured in WS (white boxes) and *clca-2* (black boxes) after 12 days of growth (A). Data are means ± SEM of 4-5 (A) or 16-20 measurements (B). For nitrate measurements, data are means ± SEM of three technical replicates of a representative experiment. Each point is a pool of at least 50 plants. Different letters at the top of the bars represent significant difference (p value<0.05).

**Supplementary Fig. S2.**
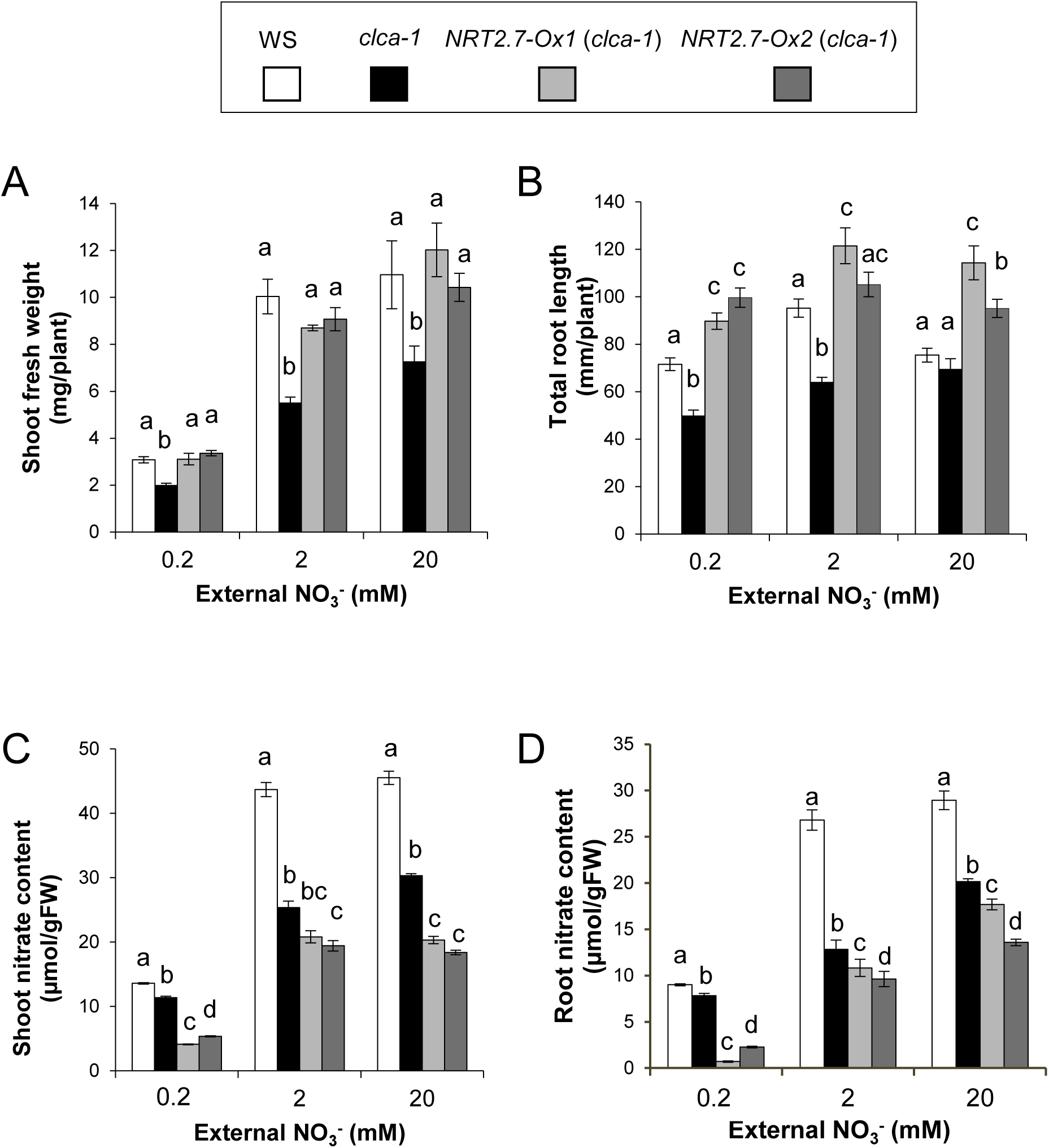
Growth and nitrate content of *NRT2.7* overexpressors in the c*lca-1* knock-out mutant backgound. Wild type (WS), *clca-1* and two over-expressing lines, *NRT2.7-Ox1* (*clca-1*) and *NRT2.7-Ox2* (*clca-1*) were grown on vertical Petri dishes for 12 days on 0.2, 2 or 20 mM external nitrate as described in the ‘Materials and Methods’. Shoot fresh weight was measured after 12 days of growth (A), and total root length were analyzed on 9 day-old plant (B). Data are averages of 4-5 (A) or 11-20 measurements (B). Shoot (C) and root (D) nitrate content in 12 day-old plants. Data are averages of three technical nitrate measurements of a representative experiment. Each point is a pool of at least 50 plants. Vertical bars represent means ± SEM. Different letters at the top of the bars represent significant difference (p value<0.05).

**Supplementary Fig. S3.**
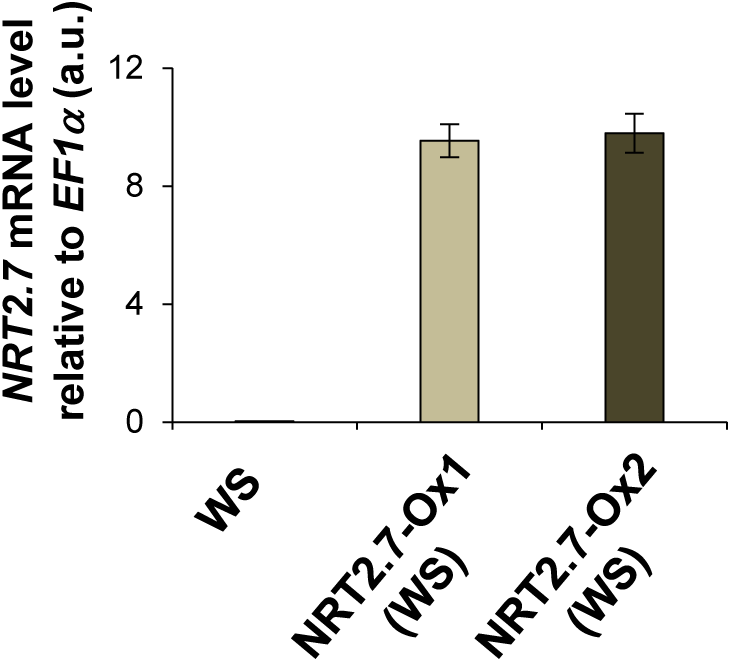
Level of *NRT2.7* expression in *NRT2.7-Ox* (WS). Plants were grown on vertical Petri dishes for 12 days on 2 mM external nitrate as described in the ‘Materials and Methods’. RT-qPCR was performed on total RNA extracted from shoot using *NRT2.7* and *EF1***α** specific primers. Data represents the means ± SEM of three biological replicates.

**Supplementary Fig. S4.**
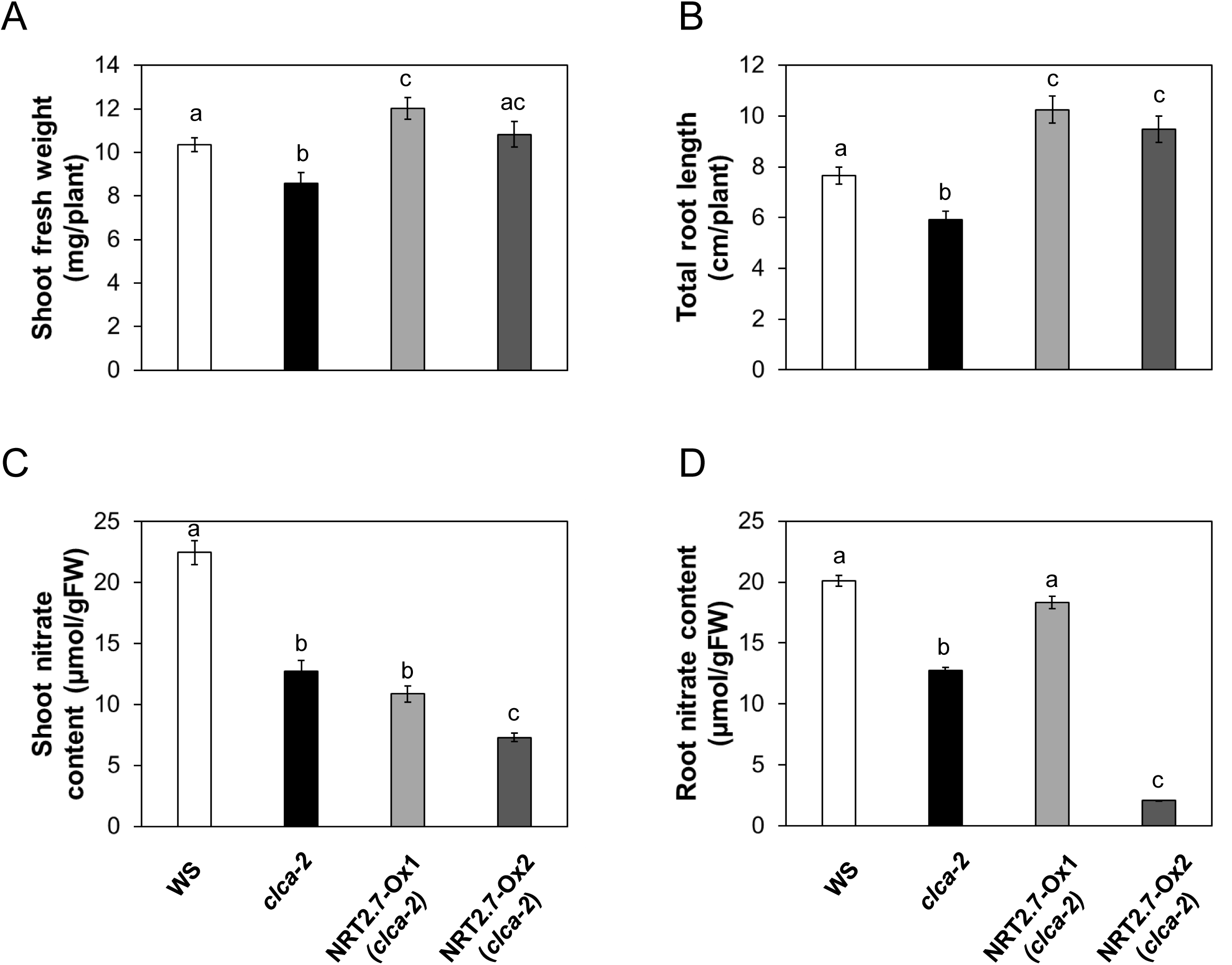
Effect of limiting N supply on the phenotype of *NRT2.7* over-expressors in *clca-2* background. Wild type (WS), *clca-2,* complemented line (*CLCa-Ox* (*clca-2*)) and *NRT2.7* over-expressing line in *clca-2* (*NRT2.7-Ox* (*clca-2*)) were grown in petri dishes for 12 days on 0.2 mM nitrate. Shoot fresh weight (A), total root length (B) and, shoot and root nitrate contents (C, D) were measured as described in Figure 1 and 2. Data are averages of three to five measurements, except for B, n=12-19 (bars representing the SEM). Different letters show significant differences between genotypes (p<0.05).

**Supplementary Fig. S5.**
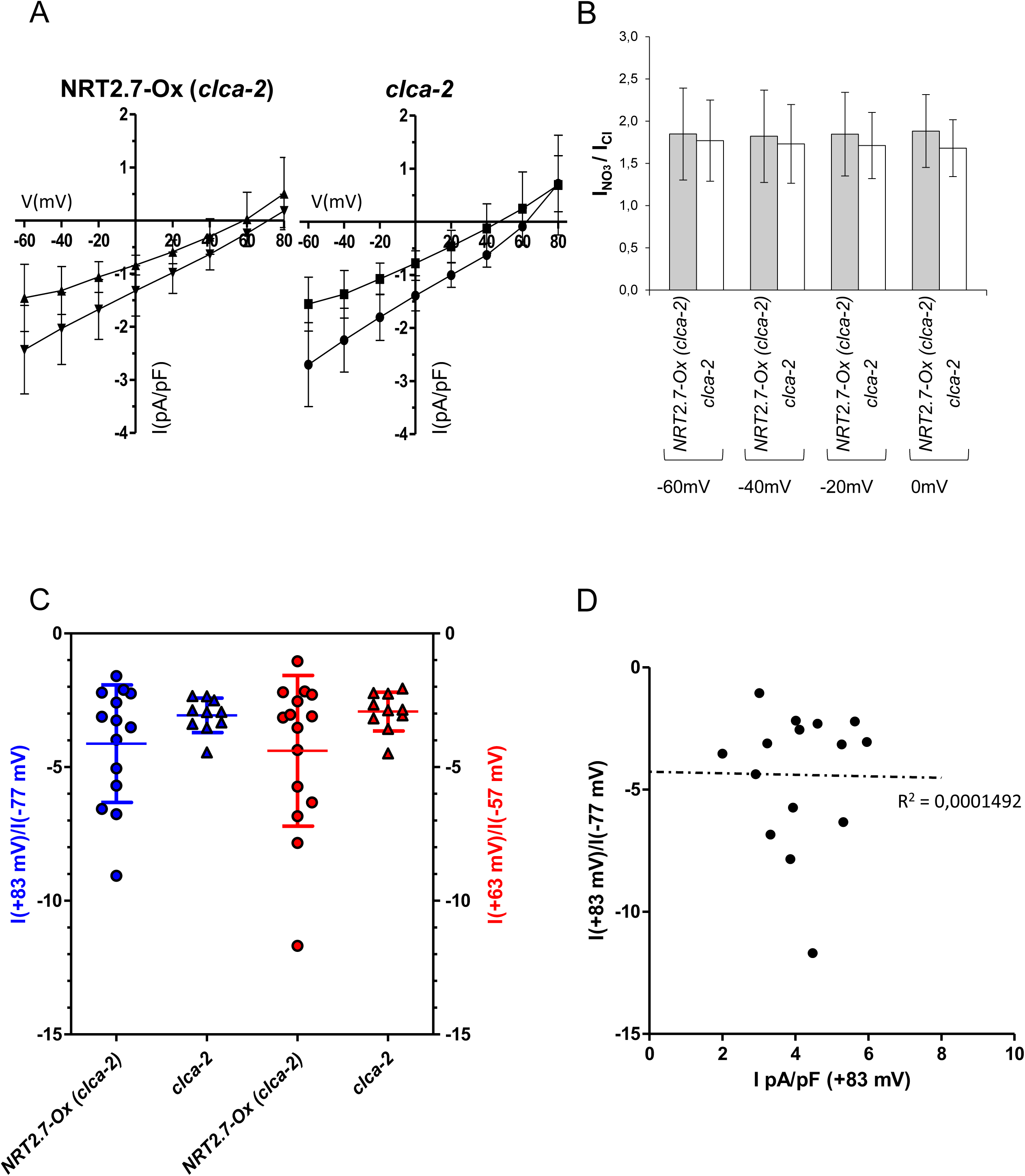
Nitrate influx is not increased in vacuoles from *NRT2.7_Ox* (*clca2*) expressing plants compared to vacuoles of *clca-2* knock-outs. (A) mean IV curves measured in vacuoles extracted from NRT2.7-Ox (*clca-2*) (top, n=6) and *clca-2* knock-out (bottom, n=6) without nitrate in the vacuolar lumen (15 mM Cl^-^, pH=5.7, see Materials and methods) and in absence (square) and presence (circle) of cytosolic nitrate (4 mM NO_3_^-^ pH=7). (B) current ratio (I_NO3_^-^/I_cl_^-^) calculated from cytosolic solution exchange experiments at different membrane potentials. Error bars are standard deviation. (C) Rectification rates calculated as I(+83mV)/I(-77mV) (blue, left axis) and I(+63mV)/I(-57mV) (red, right axis). The Rectification rate was calculated from the individual vacuoles extracted from *clca-2* knock-out mutant (triangles) and *NRT2.7-Ox* (*clca-2*) transgenic line (circles) shown in Fig. 5. Ionic currents were recorded in presence of nitrate only in the vacuolar lumen (200 mM NO_3_^-^ pH 5.5 in the vacuolar buffer, 19.2 mM Cl^-^ in the cytosolic buffer, see Materials and Methods). Statistical significance between *clca-2* knock-out mutant (triangles) and *NRT2.7-Ox* (*clca-2*) transgenic line (circles) was calculated with a non-parametrical Mann-Whitney test. (D) Plot showing the rectification rate I(+83mV)/I(-77mV) (blue, left axis) vs the current density of each vacuole of the *NRT2.7-Ox* (*clca-2*) transgenic line. The broken line shows a liner regression fit of the data illustrating the absence of correlation between the current intensity recorded in each vacuole and the rectification.

## Notes

### Competing Interest Statement

The authors have declared no competing interest.

### Summary of Updates

Figures revised: data not normalized anymore Supplemental figure S4 added. Supplemental figure S5 revised. Presisions added throughout the text.

